# Host E3 ubiquitin ligase ITCH mediates *Toxoplasma gondii* effector GRA35-triggered NLRP1 inflammasome activation and cell-autonomous immunity

**DOI:** 10.1101/2023.12.13.571530

**Authors:** Yifan Wang, L. Robert Hollingsworth, Lamba Omar Sangaré, Tatiana C. Paredes-Santos, Shruthi Krishnamurthy, Bennett H. Penn, Hao Wu, Jeroen P. J. Saeij

## Abstract

*Toxoplasma gondii* is an intracellular parasite that can activate the NLRP1 inflammasome leading to macrophage pyroptosis in Lewis rats, but the underlying mechanism is not well understood. In this study, we performed a genome-wide CRISPR screen and identified the dense granule proteins GRA35, GRA42, and GRA43 as the *Toxoplasma* effectors mediating cell death in Lewis rat macrophages. GRA35 localizes on the parasitophorous vacuole membrane, where it interacts with the host E3 ubiquitin ligase ITCH. Inhibition of proteasome activity or ITCH knockout prevented pyroptosis in *Toxoplasma*-infected Lewis rat macrophages, consistent with the “NLRP1 functional degradation model”. However, there was no evidence that ITCH directly ubiquitinates or interacts with rat NLRP1. We also found that GRA35-ITCH interaction affected *Toxoplasma* fitness in IFNγ-activated human fibroblasts, likely due to ITCH’s role in recruiting ubiquitin and the parasite-restriction factor RNF213 to the parasitophorous vacuole membrane. These findings identify a new role of host E3 ubiquitin ligase ITCH in mediating effector-triggered immunity, a critical concept that involves recognizing intracellular pathogens and initiating host innate immune responses.

**IMPORTANCE:** Effector-triggered immunity represents an innate immune defense mechanism that plays a crucial role in sensing and controlling intracellular pathogen infection. The NLRP1 inflammasome in the Lewis rats can detect *Toxoplasma* infection, which triggers proptosis in infected macrophages and eliminates the parasite’s replication niche. The work reported here revealed that host E3 ubiquitin ligase ITCH is able to recognize and interact with *Toxoplasma* effector protein GRA35 localized on the parasite-host interface, leading to NLRP1 inflammasome activation in Lewis rat macrophages. Furthermore, ITCH-GRA35 interaction contributes to the restriction of *Toxoplasma* in human fibroblasts stimulated by IFNγ. Thus, this research provides valuable insights into understanding pathogen recognition and restriction mediated by host E3 ubiquitin ligase.

## INTRODUCTION

The innate immunity system is the body’s first line of defense against infections, with cells of the innate immune system constantly recognizing infections through pattern recognition receptors (PRRs) and coordinating cellular and molecular mechanisms to mount effective antimicrobial responses. In response to particular pathogens, mammalian cells possess a sophisticated recognition mechanism named effector-triggered immunity (1). Effector-triggered immunity occurs when certain intracellular PRRs, known as Nod-like Receptors (NLRs), sense specific effectors secreted during the infection of pathogenic microbes or the alterations they induce after breaching host cell barriers. Upon detection, these NLRs can assemble into multiprotein complexes referred to as inflammasomes, which can be found in various immune cells and play critical roles in initiating host defense against infections (2). Upon recognition of pathogen effectors by inflammasome sensors, inflammatory caspases (Caspase-1, -4, or -11) are recruited and activated, leading to the release of IL1β and IL18 from infected cells (3), and inducing a form of programmed cell death known as pyroptosis via the cleavage and activation of a pore-forming protein called Gasdermin D (4, 5). Thus, the activation of inflammasomes primarily contributes to the rapid elimination of invading pathogens and is central for the mammalian innate immunity system in triggering inflammation and engaging the adaptive immune system for a more precise response.

The nucleotide-binding domain, leucine-rich repeat-containing proteins family, pyrin domain containing 1 (NLRP1) is the first NLR discovered to form an inflammasome (3). The NLRP1 inflammasome is activated by various pathogen effectors through a mechanism of “functional degradation” (6–8). The NLRP1 protein undergoes autoproteolytic processing within its function-to-find (FIIND) domain (9, 10), resulting in two polypeptides (N-terminal and C-terminal) that remain associated in an autoinhibited state. The autoinhibitory N-terminal NLRP1 polypeptide can be ubiquitinated by pathogen E3 ubiquitin ligases, such as *Shigella flexneri* IpaH7.8 (6), or processed by other pathogen proteases (e.g., *Bacillus anthracis* lethal factor and enteroviral 3C protease), leading to its ubiquitination by host N-end rule E3 ubiquitin ligases (6–8, 11, 12). This allows the active C-terminal NLRP1 polypeptide to dissociate upon proteasomal degradation of the N-terminal polypeptide and subsequently recruit Caspase-1 for inflammasome activation^12–14^. This activation process indicates that NLRP1 generally acts as a guard to sense the specific activity induced by pathogen effectors.

*Toxoplasma gondii* is an obligated intracellular pathogen and a highly successful parasite that can infect any nucleated cell and causes lifelong chronic infections in almost all warm-blooded animals. In humans, *Toxoplasma* can cause congenital infections and opportunistic infections in immunocompromised individuals (13). Although *Toxoplasma* possesses an extraordinary host range, the Lewis (LEW) rat is the only known warm-blooded animal with sterilizing immunity against the parasite (14). Polymorphisms in the rat *Nlrp1* gene determine rat strain differences in susceptibility to *Toxoplasma.* The parasite specifically activates the LEW rat NLRP1 inflammasome, resulting in macrophage pyroptosis and subsequent clearance of the infection (15–17). Moreover, NLRP1 plays a role in human monocyte control of *Toxoplasma*, and polymorphisms in *NLRP1* also influence the severity of congenital toxoplasmosis (18). However, the exact mechanism by which *Toxoplasma* activates the NLRP1 inflammasome remains unknown.

The key to the parasite’s successful survival and proliferation in diverse host cell microenvironment is that *Toxoplasma* resides within a non-fusogenic replication niche called the parasitophorous vacuole (PV), which is separated from the host cell cytoplasm by the PV membrane (PVM). Once the PV is formed, *Toxoplasma* constitutively secretes proteins from its unique organelle, dense granules, into the PV lumen (19). Many dense granule proteins (GRAs) are parasite effector proteins, which are associated with the PVM and involved in organizing the structure and environment of the PV (20), nutrient acquisition (21, 22), and modulation of host immune responses (23–26). Although most GRAs contribute to *Toxoplasma* fitness inside host cells, some of the effector proteins on the PVM can also trigger the host immunity against the parasite. For example, PVM-localized *Toxoplasma* effector GRA15 can recruit ubiquitin ligase TRAF6 to the PVM, leading to lysosomal degradation of the PV in interferon-gamma (IFNγ)-activated human fibroblast (27). Our previous study also identified three PVM-localized GRAs that are required for NLRP1 inflammasome activation in the LEW rat macrophages (28). Given that these GRA effectors were identified in single parasite clones generated from a chemical mutagenesis screen, it remains unclear whether other *Toxoplasma* effectors can trigger NLRP1 inflammasome activation as well as the mechanism involved in the inflammasome activation.

## RESULTS

### Genome-wide CRISPR screens identify *Toxoplasma* effectors that activate NLRP1 inflammasome in LEW rat macrophages

To ensure that we did not miss additional parasite effectors involved in NLRP1 inflammasome activation, we performed a genome-wide CRISPR screen in *Toxoplasma* followed by infection of LEW rat bone marrow-derived macrophages (BMDMs) (**Fig 1A**). In two independent screens, we observed a decrease in cell death of LEW rat BMDMs (**Fig 1B**) and an increase in the number of parasites that could replicate within the BMDMs (**Fig 1C**), indicating that mutant parasites that failed to activate the NLRP1 inflammasome were enriched in the population. Further selections of the parasite population allowed us to enrich for these mutant parasites, resulting in a final population where > 60% of LEW rat BMDMs were viable after infection (**Fig 1D**) and ∼ 80% of the parasites were able to replicate within the macrophages (**Fig 1C**). Using Illumina sequencing, we determined the abundance of single guide RNAs (sgRNAs) present in these parasite populations to identify enriched sgRNAs that targeted genes responsible for activating the NLRP1 inflammasome. Consistent with our previous chemical mutagenesis screen (28), our CRISPR screen enriched for only three signal peptide-coding genes, which encoded dense granule proteins GRA35, GRA42, and GRA43 (**Fig 1D** and **Table S1**). This strongly suggests that these three GRAs are the only *Toxoplasma* secreted proteins responsible for inducing pyroptosis in LEW rat BMDMs. Since we previously showed that GRA42 and GRA43 localize inside the PV lumen and facilitate the correct localization of GRA35 to the PVM (28), we focused on GRA35 hereafter to understand how this PVM-localized effector triggers NLRP1 inflammasome activation.

**Fig. 1.**
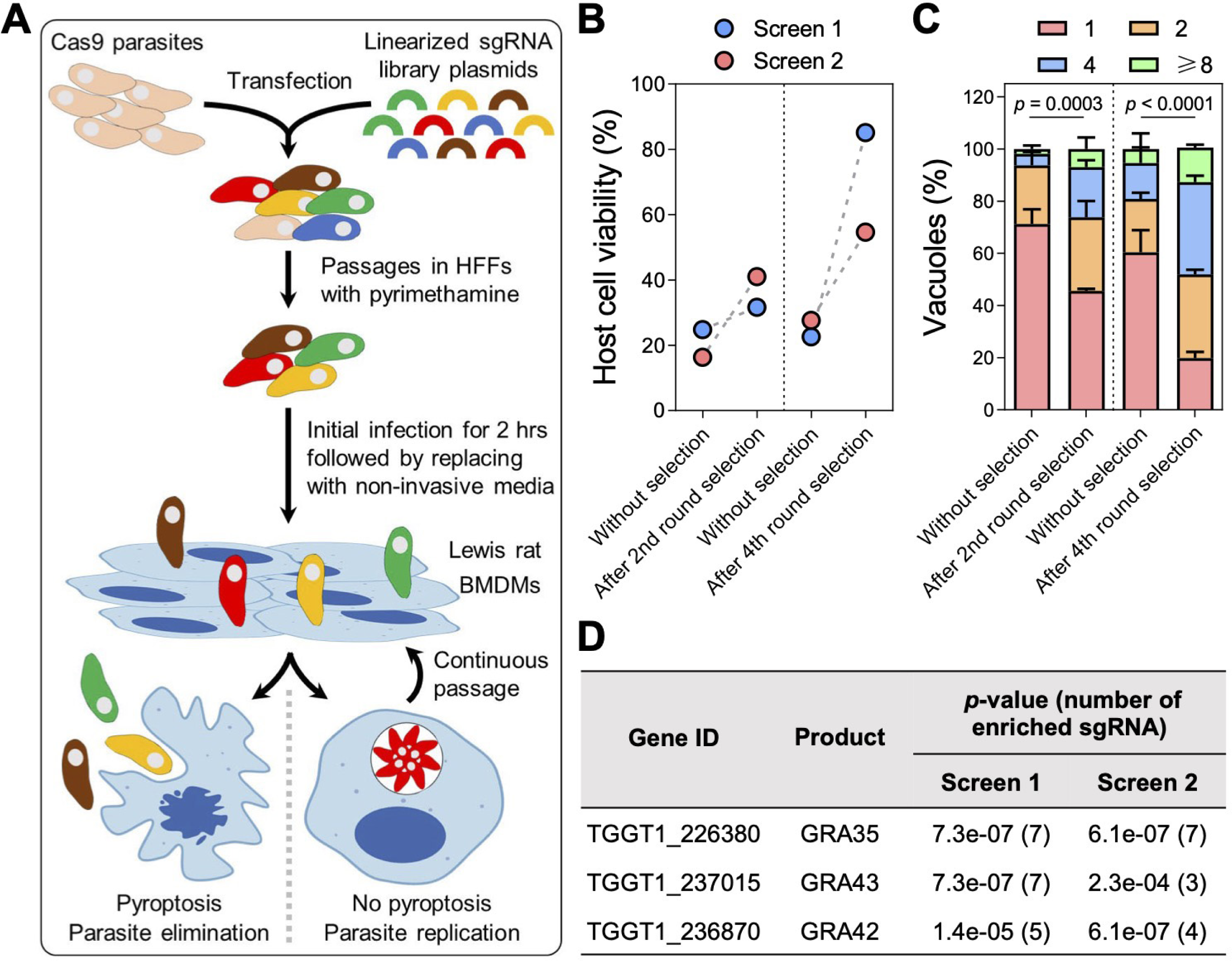
Genome-wide CRISPR screens identify *Toxoplasma* secretory effectors that induce pyroptosis in Lewis rat bone marrow-derived macrophages. **A.** Schematic of CRISPR screen. BMDMs, bone marrow-derived macrophages. **B.** Lewis rat BMDMs were infected with indicated parasites (MOI = 1) for 24 h. Macrophage viability was measured via MTS assay. Data are displayed as paired plots for each individual screen. **C.** Lewis rat BMDMs were infected with indicated parasites (MOI = 0.5) for 24 h. The number of parasites per vacuole was quantified by microscopy. A total of 100 to 120 vacuoles were counted per experiment. Data are displayed as mean + SD with two independent experiments. Significance was determined with two-way ANOVA with Tukey’s multiple comparisons test. **D.** Top candidate genes with at least 3 enriched sgRNAs and significant enrichment (*p*-value < 0.05, analyzed by MAGeCK algorithm) after 4th round selection in both screens. Numbers between parentheses are the number of enriched sgRNAs after the 4^th^ round of selection.

### GRA35 interacts with host E3 ubiquitin ligase ITCH

GRA35 is not predicted to have proteolytic or ubiquitin ligase activity. Therefore, to gain insight into the molecular mechanism underlying GRA35-mediated NLRP1 inflammasome activation, we first determined the topology of GRA35 on the PVM. GRA35 has a single transmembrane domain that separates the protein into a short N-terminus (97 amino acids) and a long C-terminus (237 amino acids). To clarify its topology, we selectively permeabilized the host plasma membrane, but not the PVM, using 0.001% Digitonin in host cells infected with a parasite strain expressing GRA35 C-terminally tagged with the HA epitope (**Fig 2A**).

**Fig. 2.**
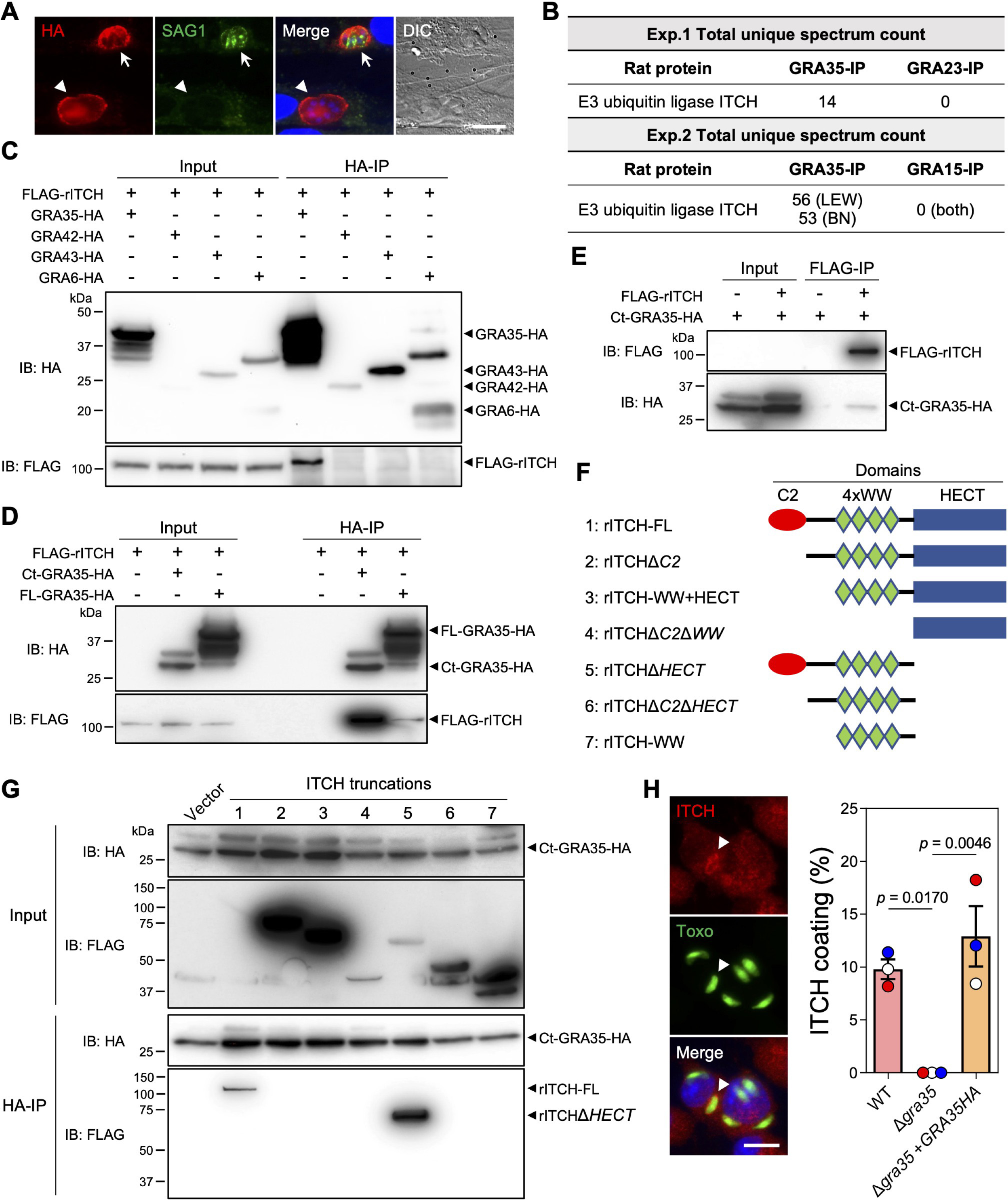
GRA35 interacts with host E3 ubiquitin ligase ITCH. **A.** HFFs were infected with parasites expressing GRA35 endogenously tagged at the C-terminus with the HA epitope for 16 h. After fixation, the cells were permeabilized with 0.001% digitonin followed by staining with antibodies against SAG1 and the HA epitope. The images are representative of results from 2 independent experiments (scale bar = 5 μm). Arrows indicate a fully permeabilized vacuole and arrowheads denote the vacuole that is not permeabilized. **B.** Lysates from Lewis rat BMDMs infected with parasites expressing C-terminal HA-tagged GRA35 or other PVM-localized proteins (GRA23 or GRA15, served as negative controls) were immunoprecipitated using HA antibodies followed by mass spectrometry analysis. The total unique spectrum count of host E3 ubiquitin ligase ITCH is indicated. **C.** Lysates of HEK293T cells transiently expressing FLAG-rITCH with the indicated GRA fused with HA epitope were immunoprecipitated using HA antibodies followed by immunoblotting (IB) analysis with the indicated antibodies. 5% of the total lysate was loaded and used as input. The images are representative of results from 2 independent experiments. **D.** HEK293T cells transiently expressing FLAG-rITCH and either full-length (FL) or the C-terminus (Ct) of GRA35 fused with the HA epitope were analyzed as in (c). The images are representative of results from 2 independent experiments. **E.** HEK293T cells transiently expressing FLAG-rITCH and an HA-tagged GRA35 C-terminal fragment were immunoprecipitated using FLAG antibodies and analyzed as in (c). The images are representative of results from 2 independent experiments. **F.** Schematic illustration of the constructs used for the generation of full-length or truncated rat ITCH. **G.** HEK293T cells transiently expressing a C-terminal fragment of GRA35 fused with the HA epitope and FLAG-tagged rat ITCH truncations (indicated in f) were immunoprecipitated using HA antibodies and analyzed as in (c). The images are representative of results from 2 independent experiments. **H.** Lewis rat BMDMs infected with GFP-expressing *Toxoplasma* for 4h were stained with antibodies against ITCH. The images are representative of results from wild-type parasite infected BMDMs (scale bar = 10 μm). The percentage of vacuoles coated with ITCH was quantified as shown on the right. Data are displayed as mean ± SD with independent experiments (*n* = 3) indicated by the same color dots. Significance was determined with one-way ANOVA with Tukey’s multiple comparisons test. Arrowhead indicates the vacuole coated with ITCH.

We used SAG1 antibodies, which stain the parasite plasma membrane, to identify fully permeabilized vacuoles in infected cells. In host cells containing SAG1-negative vacuoles, we observed HA antibody staining on the PVM (**Fig 2A**), indicating that GRA35 was localized on the PVM with its C-terminus facing the host cytosol. The C-terminus of GRA35 contains several coiled-coil domains (28), which are often involved in protein-protein interactions. However, we previously did not observe a direct interaction between GRA35 and LEW rat NLRP1 (28), indicating that *Toxoplasma* GRA35 likely induces NLRP1 inflammasome activation by interacting with other host proteins. To identify GRA35 interaction partners, we immunoprecipitated GRA35 from *Toxoplasma*-infected rat BMDMs. Mass spectrometry analysis identified only one rat protein, the E3 ubiquitin ligase ITCH, that was specifically and consistently present in GRA35 immunoprecipitated samples (**Fig 2B**). To confirm the direct interaction between GRA35 and rat ITCH, we performed coimmunoprecipitation experiments in HEK293T cells expressing FLAG-tagged rat ITCH (FLAG-rITCH) and HA-tagged GRA35 or HA-tagged control dense granule proteins (e.g., GRA42, GRA43, and GRA6) (**Fig 2C**). We observed that GRA35, but not the control dense granule proteins, specifically immunoprecipitated rat ITCH (**Fig 2C**). Furthermore, we found that the C-terminus (amino acid 142 to 378) of GRA35 had a stronger affinity for rat ITCH than full-length GRA35 (**Fig 2D**), indicating that the C-terminus serves as the functional domain of GRA35 for interacting with host proteins. To confirm the direct interaction between rat ITCH and *Toxoplasma* GRA35, we performed reverse immunoprecipitations and found that rat ITCH binds specifically to the C-terminus of GRA35 (**Fig 2E**). ITCH is a member of the NEDD4 family of E3 ubiquitin ligases and has several domains, including an N-terminal Ca^2+^-dependent phospholipid-binding C2 domain, four tandem WW domains for substrate binding, and the C-terminal HECT domain for interaction with an E2 ubiquitin transferase, leading to the ubiquitination of substrates(29). To determine the domain of ITCH that binds to *Toxoplasma* GRA35, we generated FLAG-tagged constructs expressing different ITCH truncations (**Fig 2F**). By performing coimmunoprecipitations with the GRA35 C-terminus, we found that ITCH only binds to GRA35 in the presence of its N-terminal C2 domain (**Fig 2G**). Given that the C2 domain of NEDD4 family E3 ubiquitin ligases mainly involves binding to membranes (30, 31), we determined the localization of ITCH in *Toxoplasma*-infected LEW rat BMDMs (**Fig 2H**). We found that ITCH is recruited to the PVM of ∼10% of wild-type parasites, whereas ITCH PVM coating is completely absent in BMDMs infected with Δ*gra35* parasites (**Fig 2H**). Taken together, these results demonstrate that the *Toxoplasma* effector GRA35 localizes on the PVM where its host-cytosol facing C-terminus recruits the host E3 ubiquitin ligase ITCH.

### ITCH mediates NLRP1 inflammasome activation triggered by *Toxoplasma* infection in LEW rat macrophages

As a HECT-type E3 ubiquitin ligase, ITCH primarily targets substrates for K48-linked ubiquitination, which is a well-established signal for canonical proteasomal degradation (32). Consistent with the observation that blocking proteasome activity prevents NLRP1 inflammasome activation (6, 33, 34), we found that *Toxoplasma* was unable to induce cell death in LEW rat BMDMs in the presence of the proteasome inhibitor MG132 (**Fig 3A**). MG132 treatment did not block parasite invasion (**Fig S1**), suggesting that *Toxoplasma*-induced LEW rat NLRP1 inflammasome activation is also mediated by the “functional degradation” of the repressive NLRP1 N-terminus. To further understand the role of ITCH in mediating NLRP1 inflammasome activation in LEW rat BMDMs after *Toxoplasma* infection, we generated *Itch* knockout BMDMs by delivering recombinant Cas9 protein in a complex with three sgRNAs targeting the second exon (only ∼70bp in the first exon) of rat *Itch* (**Fig 3B**). PCR amplification of a region containing the *Itch* sgRNA targeting sites resulted in reduced band size compared to negative control cells (**Fig 3C**), and Sanger sequencing confirmed that the *Itch* locus is disrupted at the sgRNA targeting sites (**Fig S2**). Inference of CRISPR Edits (ICE) analysis of the sequencing products revealed that 86% ± 4.1% of cells (n = 4) had CRISPR editing in the *Itch* locus around the sgRNA targeting sites (**Fig 3D**). As a positive control, we also generated *Nlrp1* knockout LEW rat BMDMs using a similar approach, which resulted in 96% ± 1.9% (n = 4) editing efficiency in the *Nlrp1* locus (**Fig 3D**). Significantly less cell death was observed in BMDMs transfected with *Itch* or *Nlrp1* sgRNAs after infection with *Toxoplasma* compared to BMDMs transfected with Cas9 protein alone or Cas9 protein with scrambled sgRNAs (targeting *E. coli* LacZ) (**Fig 3E**). Additionally, there was increased parasite replication in BMDMs transfected with *Itch* or *Nlrp1* sgRNAs, indicated by a significantly higher proportion of vacuoles containing more than 1 parasite compared to control BMDMs (**Fig 3E**). Taken together, these results reveal that the host E3 ubiquitin ligase ITCH plays an important role in the NLRP1-mediated cell death upon *Toxoplasma* infection of LEW rat BMDMs.

**Fig. 3.**
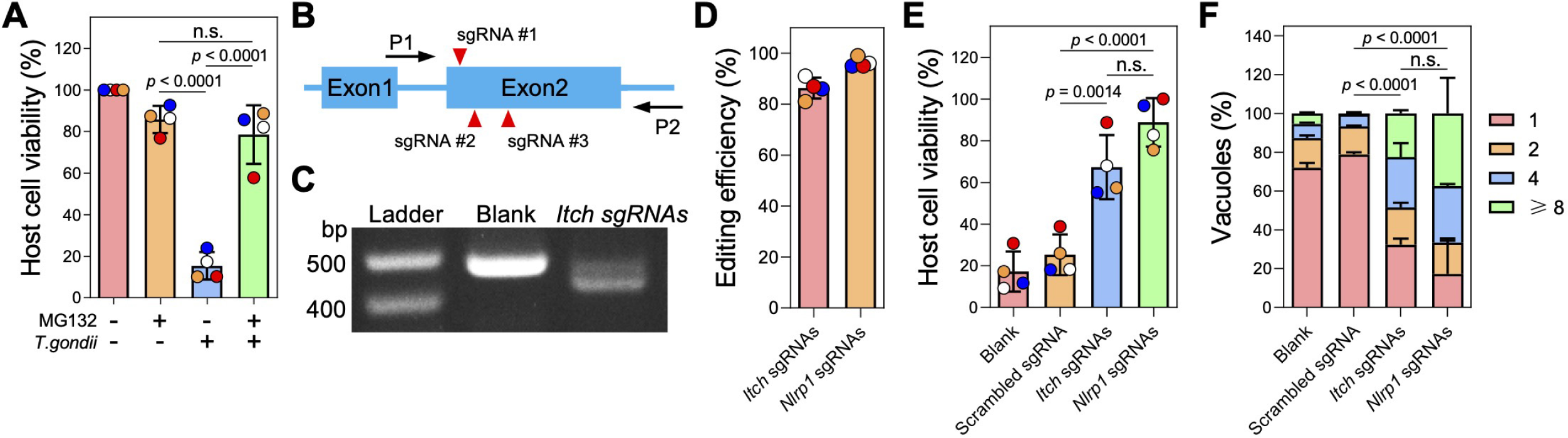
Knockout of *Itch* impairs *Toxoplasma*-induced NLRP1-mediated cell death of Lewis rat BMDMs. **A.** Lewis rat BMDMs pre-treated with 0.5 μM proteasome inhibitor MG132 or left untreated for 2 h were infected with *Toxoplasma* parasites (MOI = 1) for 24 h. Macrophage viability was measured via MTS assay. Data are displayed as mean ± SD with independent experiments (*n* = 4) indicated by the same color dots. Significance was determined with one-way ANOVA with Tukey’s multiple comparisons test. **B.** Schematic illustration of sgRNA targeting sites in the first two exons of rat *ITCH* locus. P1 and P2 are the primers used for amplifying the sgRNA targeting region and verification by Sanger sequencing. **C.** PCR amplification of the *Itch* sgRNA targeting region using P1 and P2 from Lewis rat BMDMs transfected with Cas9 protein only (blank) or Cas9 protein assembled with Itch sgRNAs. **D.** Lewis rat BMDMs were transfected with *in vitro* assembled CRISPR/Cas9 ribonucleoprotein containing indicated sgRNAs. The editing efficiency of *Itch* or *Nlrp1* was analyzed from the Sanger sequencing products (shown in c) using the Inference of CRISPR Edits (ICE) online tool (https://www.synthego.com/products/bioinformatics/crispr-analysis). The percentage of CRISPR edited *Itch* or *Nlrp1* is displayed as mean ± SD with independent experiments (*n* = 4) indicated by the same color dots. **E.** Lewis rat BMDMs generated in (d) were infected with wild-type *Toxoplasma* (MOI = 1) for 24 h. Macrophage viability was measured via MTS assay. Data are displayed as mean ± SD with independent experiments (*n* = 4) indicated by the same color dots. Significance was determined with one-way ANOVA with Tukey’s multiple comparisons test. **F.** Lewis rat BMDMs transfected with *in vitro* assembled CRISPR/Cas9 ribonucleoprotein containing indicated sgRNAs were infected with wild-type *Toxoplasma* (MOI = 0.5) for 24 h. The number of parasites per vacuole was quantified by microscopy. A total of 100 to 120 vacuoles were counted per experiment. Data are displayed as mean + SD with 2 independent experiments. Significance was determined with two-way ANOVA with Tukey’s multiple comparisons test.

To determine if ITCH directly ubiquitinates the LEW rat NLRP1, we performed an *in vitro* ubiquitination assay using recombinant human ITCH protein (GRA35 also interacts with human ITCH as shown in **Fig 4A**), which shares over 90% homology with rat ITCH, and FLAG-tagged LEW rat NLRP1 protein produced from SF9 insect cells (**Fig S3A**). However, we only observed a strong auto-ubiquitination of ITCH in the presence of ATP and did not detect any ubiquitination of NLRP1. While this assay allowed us to assess the direct ubiquitination of NLRP1 by ITCH *in vitro*, it remained unclear whether *Toxoplasma* infection triggers the ubiquitination activity of ITCH, particularly in the context of ITCH interacting with GRA35. To address this question, we performed an in-cell ubiquitination assay in HEK293T cells that stably expressed LEW rat NLRP1 fused with EGFP at its N-terminus and tagged with the MYC epitope at its C-terminus. We co-expressed HA-tagged ubiquitin and FLAG-tagged ITCH into the cells and then performed GFP-immunoprecipitation to capture the NLRP1 protein. However, we did not observe any ubiquitination of NLRP1 regardless of *Toxoplasma* infection (**Fig S3B**). We also found that ITCH did not directly interact with rat NLRP1 in the same experiment (**Fig S3B**).

**Fig. 4.**
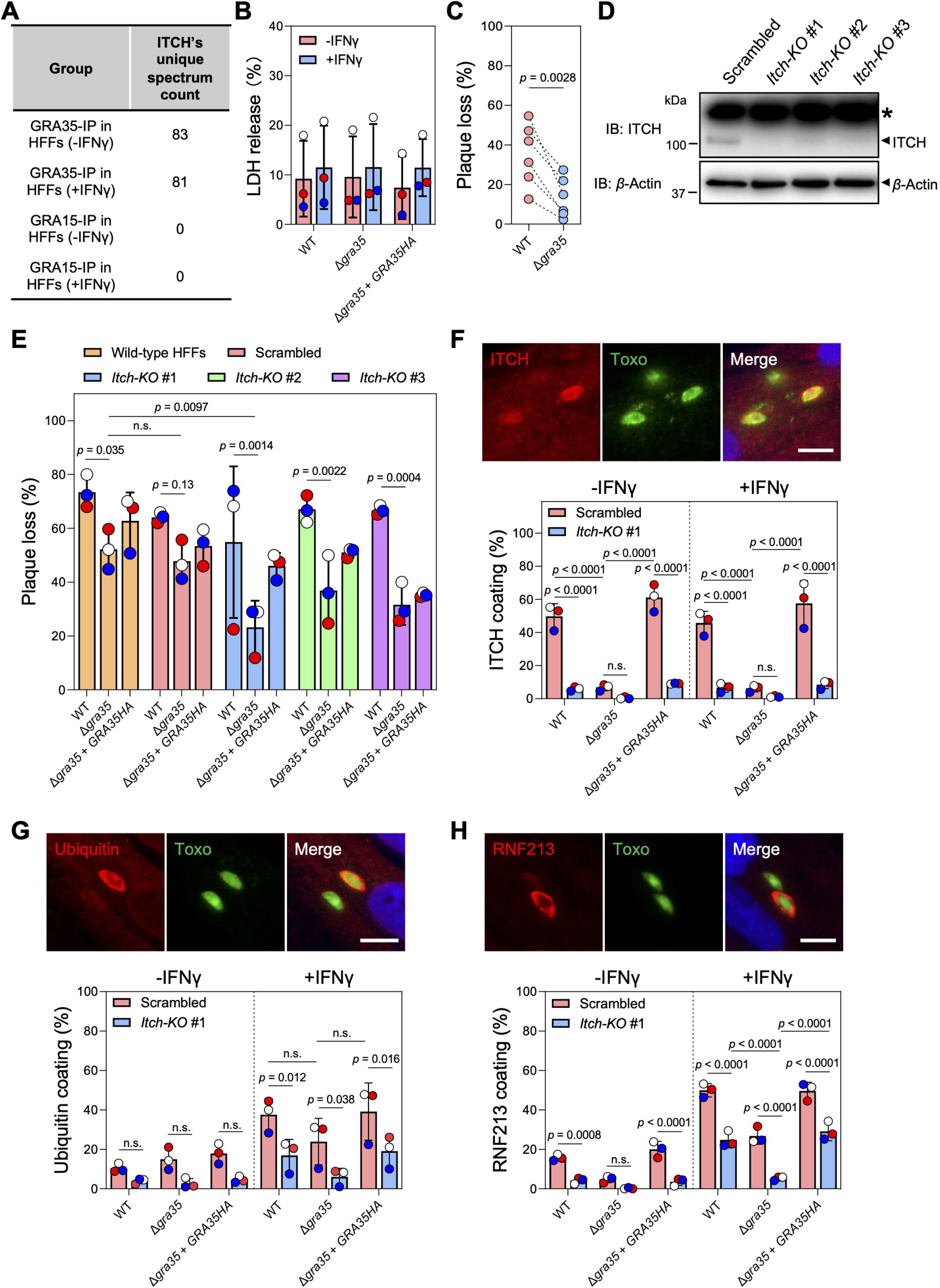
ITCH impacts ubiquitin and RNF213 recruitment to the PVM and alters the cell-autonomous response to *Toxoplasma* infection in human fibroblasts. **A.** ITCH was identified as interacting with GRA35 in our immunoprecipitation coupled with mass spectrometry analysis in naïve and IFNγ-activated HFFs^28^. The total unique spectrum count of human ITCH identified in GRA35-immunoprecipitated samples is shown. GRA15 serves as a PVM-localized control. **B.** HFFs pre-stimulated with 10 U/ml IFNγ or unstimulated for 24 h were infected with indicated parasite strains (MOI = 1) for another 24 h. Cell viability was assessed via LDH assay. Data are displayed as mean ± SD with independent experiments (*n* = 3) shown by the same color dots. Significance was determined with two-way ANOVA with Tukey’s multiple comparisons test. **C.** HFFs pre-stimulated with 10 U/ml IFNγ or left unstimulated for 24 h were infected with indicated parasite strains (100 parasites per well). Plaque number was measured at 5 days post-infection. The loss of plaques in IFNγ-stimulated HFFs was calculated relative to unstimulated HFFs. Data are displayed as paired scatterplots, and significance was determined with two-tailed paired *t*-test. **D.** HFFs were transduced with lentivirus containing 3 individual ITCH-targeting or scrambled sgRNAs followed by puromycin selection. Lysates of these HFFs were analyzed by immunoblotting with ITCH and Actin antibodies. The images are representative of results from 2 independent experiments. Star (*) indicates a non-specific band. **E.** Indicated HFFs pre-stimulated with 10 U/ml IFNγ or left unstimulated for 24 h were infected with WT, Δ*gra35*, or Δ*gra35* + *GRA35HA* strains (100 parasites per well). Plaque numbers were counted 5 days post-infection. The loss of plaques in IFNγ-stimulated HFFs was calculated and expressed relative to unstimulated HFFs. Data are displayed as mean ± SD with independent experiments (*n* = 3) indicated by the same color dots. Significance was determined with two-way ANOVA with Tukey’s multiple comparisons test. **F-H.** Indicated HFFs pre-stimulated with 10 U/ml IFNγ or left unstimulated for 24 h were infected with parasites (MOI = 1) for 4 h. The cells were fixed and stained for ITCH (**F**), Ubiquitin (**G**), or RNF213 (**H**). Representative images from wild-type parasite infection in IFNγ-activated control HFFs (transduced with scrambled sgRNAs) are shown (scale bar = 10 μm). The percentage of vacuoles coated with ITCH, Ubiquitin, or RNF213 was quantified. Data are displayed as mean ± SD with independent experiments (*n* = 3) shown by colored dots. Significance was determined with two-way ANOVA with Tukey’s multiple comparisons test.

Previous studies have shown that the host serine dipeptidase DPP9 can inhibit NLRP1 inflammasome activation by sequestering the free NLRP1 C-terminus and thereby blocking NLRP1 assembly with Caspase-1 (35, 36). The inhibition of DPP9 with a small molecule called Val-boroPro (VbP) causes NLRP1 inflammasome activation in various cell types (33, 34, 37). VbP treatment specifically induced cell death in BMDMs isolated from LEW rats but not Brown Norway (BN) rats, which have an NLRP1 inflammasome that cannot recognize *Toxoplasma* infection (**Fig S4A**). This suggests that *Toxoplasma* may activate the NLRP1 inflammasome via acting on DPP9. VbP treatment induced cell death in control and *Itch* knockout rat BMDMs but not in *Nlrp1*-knockout BMDMs (**Fig S4B**). To investigate whether ITCH acts upstream of DPP9 to activate the NLRP1 inflammasome, we determined if ITCH interacts with DPP9 and mediates its ubiquitination. However, we found that ITCH neither directly ubiquitinates DPP9 *in vitro* (**Fig S3A**) nor interacts with DPP9 (**Fig S4C**). Surprisingly, our previous transcriptomic analysis (15) of the BMDMs isolated from various rat strains indicated that DPP9 is not even expressed in rat macrophages (**Fig S4D**). Collectively, these results suggest that ITCH mediates NLRP1 inflammasome activation in *Toxoplasma*-infected LEW rat macrophages using a yet-to-be-determined mechanism that is independent of NLRP1 ubiquitination and DPP9 inhibition.

### ITCH-GRA35 interaction affects *Toxoplasma* susceptibility to IFNγ-induced growth inhibition in human fibroblasts

ITCH plays a vital role in immune-related functions, such as T-cell responses and apoptosis, across a range of host species (38). Moreover, our previous proteomic analysis in human foreskin fibroblasts (HFFs) (27) indicated that GRA35 also interacts with human ITCH in the presence or absence of IFNγ activation (**Fig 4A** and **Table S3**). Therefore, we sought to determine whether GRA35-ITCH interaction triggers host mechanisms response to *Toxoplasma* infection other than inflammasome activation in rat macrophages. Firstly, we investigated whether GRA35 plays a role in pyroptotic cell death in IFNγ-activated HFFs. To do so, we infected both naive and IFNγ-activated HFFs with wild-type, GRA35-complemented, and Δ*gra35* parasites. We found that the levels of LDH release, a marker of host cell death, were similar between HFFs infected with wild-type, Δ*gra35*, and GRA35-complemented parasites (**Fig 4B**). While the overall levels of cell death did not change significantly, the absence of GRA35 led to significantly lower growth inhibition in IFNγ-activated HFFs (**Fig 4C**). This indicates that GRA35 is involved in determining *Toxoplasma* susceptibility to IFNγ-mediated restriction in HFFs.

To determine whether ITCH is involved in IFNγ-induced parasite inhibition, we used the CRISPR/Cas9 technique to generate pooled *Itch*-knockout HFFs (**Fig 4D**). ITCH knockout did not affect the growth of wild-type parasites in the presence of IFNγ but caused Δ*gra35* parasites to become significantly more resistant to IFNγ-induced growth inhibition in ITCH knockout HFFs (**Fig 4E**). This result indicates that ITCH contributes to IFNγ-mediated parasite restriction. When cells are activated by IFNγ, the pathogen-containing vacuole membrane of several intracellular pathogens, including *Toxoplasma*, can be marked with polyubiquitin chains, which initiates a cascade of molecular events leading to vacuole disruption (39). To understand how the interaction between GRA35 and ITCH affects IFNγ-induced parasite growth inhibition, we examined the recruitment of ITCH (**Fig 4F**) and ubiquitin (**Fig 4G**) to the PVM in control (scrambled sgRNA-transduced) and *Itch*-knockout HFFs. We found that 50∼60% of vacuoles of wild-type or GRA35-complemented parasites were coated with ITCH in control HFFs, regardless of IFNγ activation, whereas Δ*gra35* parasites had significantly lower ITCH recruitment (**Fig 4F**). However, a significantly lower level of ubiquitin coating was observed in *Itch*-knockout HFFs in the presence of IFNγ compared to without IFNγ activation regardless of GRA35 expression (**Fig 4G**). These results indicate that ITCH does not exclusively mediate the ubiquitination of the vacuole in HFFs and may affect the dynamics of other host E3 ubiquitin ligases that mark the PVM with ubiquitin. RNF213, another host E3 ubiquitin ligase that is an IFNγ-stimulated gene and actively accumulates on the vacuole, mediates *Toxoplasma* growth inhibition (40, 41). We found that Δ*gra35* parasites had significantly less RNF213 on the PVM, especially in IFNγ-activated *Itch*-knockout HFFs, compared to wild-type and Δ*gra35* + *GRA35HA* parasites (**Fig 4H**). Taken together, these results indicate that the interaction between ITCH and GRA35 plays a role in determining parasite fitness in human cells by affecting the RNF213 loading and ubiquitin status on the vacuole.

## DISCUSSION

The intricate interplay involving the *Toxoplasma* infection and the activation of effector-triggered immunity provides profound insight into the complexity of host-pathogen interactions and the robustness of the mammalian immune system. This study demonstrates that the *Toxoplasma* secreted effector GRA35 can interact with the host E3 ubiquitin ligase ITCH, which recruits this ubiquitin ligase to the parasite’s replication niche. E3 ubiquitin ligases are a group of critical enzymes in the host cell’s ubiquitination system, contributing significantly to various cellular processes, including protein degradation and turnover, cell cycle progression, and signal transduction (42). Besides maintaining the homeostasis of host cells, E3 ubiquitin ligases also participate in host immune responses. Particularly, our results reveal that E3 ubiquitin ligase ITCH mediates the recognition of *Toxoplasma* infection via its interaction with GRA35, which leads to NLRP1 inflammasome activation in rat macrophages and cell-autonomous response in human fibroblast activated by IFNγ. Eventually, both pathways contribute to parasite restriction in the host cells, highlighting the novel role of E3 ubiquitin ligase ITCH in mediating effector-triggered immunity and acting as a sentinel against pathogen infection across host species and cell types.

Although the exact molecular mechanism of GRA35-ITCH interaction-induced inflammasome activation is still unclear, inhibition of proteasome activity blocks pyroptosis in *Toxoplasma*-infected macrophages in a manner consistent with the “functional degradation” model for NLRP1 activation. However, we found that ITCH does not directly ubiquitinate or interact with LEW rat NLRP1, suggesting that ITCH may act on another host protein to initiate the “functional degradation” of NLRP1. Given that ITCH is not involved in DPP9 inhibition, another E3 ubiquitin ligase may be the potential host protein that is directly involved in NLRP1 inflammasome activation. It was also intriguing to discover that DPP9 is not expressed in LEW rat macrophages, but VbP still efficiently causes NLRP1 inflammasome activation. This implies that VbP not only targets DPP9 but also other host proteins that could maintain the homeostasis of the NLRP1 inflammasome. On the other hand, our results suggest that the mechanism of NLRP1 inflammasome activation induced by VbP and *Toxoplasma* is different, and that the interaction between ITCH and GRA35 on the PVM is a unique mechanism for the host to recognize *Toxoplasma* infection. Our data indicate that ITCH interacts with GRA35 via its C2 domain, which is known for membrane binding as well as mediating protein oligomerization (43). Thus, one possibility is that GRA35 could mediate ITCH oligomerization, which further enhances the ubiquitin ligase activity (44) and promotes its binding to the substrates, potentially including proteins that modulate LEW rat NLRP1 stability. Alternatively, this binding event could initiate broader perturbations to host cell homeostasis that are ultimately integrated through the inflammasome response.

In previous proteomic studies, ITCH was also identified as a host protein that is enriched on the vacuole in parasite-infected HFFs (45, 46). Our study shows that in addition to its role in mediating NLRP1 inflammasome activation in LEW rat macrophages, ITCH also contributes to IFNγ-mediated parasite restriction in human fibroblasts. When human and murine cells are activated by IFNγ, vacuole ubiquitination occurs, leading to vacuole disruption and parasite inhibition (39). However, only a few host E3 ubiquitin ligases have been found to target the vacuole and participate in its ubiquitination. Although ITCH recruitment to the vacuole is solely mediated by GRA35 and is not induced by IFNγ in HFFs, the deletion of ITCH affects the loading of ubiquitin and the newly identified parasite restriction factor, RNF213, to the vacuole. As a result, less growth inhibition was observed for Δ*gra35* parasites in *Itch* knockout HFFs (**Fig 4E**). Notably, the loading of RNF213 on the vacuole of wild-type parasites is also decreased in *Itch*-knockout HFFs, but parasite growth inhibition remains similar to control HFFs, indicating that other host IFNγ-stimulated genes are involved in this process and may target the PVM in a GRA35 dependent manner. We also found that ITCH coating in Δ*gra35* parasites was completely abolished in LEW rat macrophages (**Fig 2H**), but not in HFFs (**Fig 4F**), suggesting that other *Toxoplasma* proteins localized on the PVM particularly involved in the recruitment of human ITCH to the PVM. Further studies are required to determine the exact mechanism underlying ITCH-mediated parasite growth inhibition and explore the interactions of other parasite proteins with ITCH from different host origins.

*Toxoplasma* has long been regarded as a successful and persistent intracellular pathogen, owing to its capacity to establish lifelong chronic infection in a wide range of warm-blooded animals. However, carrying effector proteins like GRA35, which can be recognized by the host innate immune machinery, appears counterproductive to *Toxoplasma*’s survival and proliferation in the host cells. This seeming contradiction highlights the intricate dynamics of host-pathogen interactions. Many effector proteins released by *Toxoplasma* during host infection play critical roles in mediating the evasion of the innate immune response and determining parasite virulence (47, 48). Nevertheless, a few “detrimental” *Toxoplasma* effectors could temper the host’s immune response to prevent parasite overgrowing, which might otherwise lead to host death, an unfavorable outcome for the parasite given it relies on the host for its propagation. Therefore, the detriment effector-triggered immunity poses to the parasite’s immediate survival could serve a greater role in ensuring long-term persistence and transmission. It is worth noticing that GRA35 might also play a role in enhancing parasite fitness in other contexts. For instance, GRA35 was recently identified as the top hit that maintains the growth and proliferation of type II *Toxoplasma* PRU strain in IFNγ-activated human fibroblasts (49), which is different from the “detrimental” role of GRA35 played in the type I RH parasites demonstrated in our current study. Given that the C-terminus of GRA35 (the part facing to host cytosol) has a high rate of nonsynonymous/synonymous (NS/S) polymorphisms among 64 different *Toxoplasma* strains (28), GRA35 has likely undergone positive selection due to the host immune pressure. Since ITCH interacts with GRA35 via its C-terminus, it would be interesting to know if the polymorphisms of GRA35 C-terminus affect the efficiency of ITCH recognition and, furthermore, influence the outcome of *Toxoplasma* infection.

Although more research is needed to fully understand the mechanistic details, host E3 ubiquitin ligases clearly represent a critical checkpoint in anti-*Toxoplasma* cell-autonomous response. In addition to ITCH and RNF213, other host E3 ubiquitin ligases were discovered to be involved in *Toxoplasma* restriction, such as TRIM21 (50) and TRAF6 (27). Addressing the precise mechanisms through which these enzymes restrict *Toxoplasma* infection could pave the way for developing innovative strategies to combat toxoplasmosis and provide novel insight into our understanding of effector-triggered immunity against other intracellular pathogens from the perspective of host-pathogen interaction.

## METHODS & MATERIALS

### Reagents and antibodies

Dextran sulfate sodium salt was purchased from Santa Cruz Biotechnology (Cat# sc-203917). CellTiter 96^®^ AQueous One Solution Cell Proliferation Assay (MTS reagent) was obtained from Promega (Cat# G3580). Proteasome inhibitor MG132 (Cat# S2619) and Caspase-1/11 inhibitor VX-765 (Cat# S2228) were purchased from Selleck Chemicals. Digitonin (Cat# 300410) and Val-boroPro (Cat# 5314650001) were obtained from Sigma-Aldrich. Halt^TM^ protease and phosphatase inhibitor cocktail (Cat # 78444) was purchased from Thermo Scientific.

Pierce^TM^ anti-HA magnetic beads (Cat# 88837) were purchased from Thermo Scientific. Rat monoclonal anti-HA (3F10) antibodies (Cat# 11867431001), Mouse monoclonal anti-FLAG (M2) antibodies (Cat# F3165), and Horseradish peroxidase (HRP)-conjugated Mouse monoclonal anti-FLAG antibodies (Cat# A8592) were purchased from Sigma-Aldrich. Rabbit monoclonal anti-FLAG (D6W5B) antibodies (Cat # 14793S) and Mouse monoclonal anti-MYC (9B11) antibodies (Cat# 2276S) were obtained from Cell Signaling Technology. Mouse polyclonal anti-V5 antibodies were purchased from MBL Life Science (Cat# PM003). Rabbit polyclonal anti-GFP antibodies were purchased from Novus Biologicals (Cat# NB600-308). Rabbit monoclonal anti-ITCH (D8Q6D) antibodies used for immunoblotting were purchased from Cell Signaling Technology (Cat# 12117S), and purified Mouse anti-ITCH antibodies used for immunofluorescence assay were purchased from BD Biosciences (Cat# 611198). Mouse monoclonal anti-Ubiquitin antibodies used for immunoblotting were obtained from Santa Cruz Biotechnology (Cat# sc-8017), and Mouse monoclonal anti-Ubiquitin antibodies used for immunofluorescence assay were purchased from Enzo Life Sciences (Cat# ENZ-ABS840-0100). Rabbit polyclonal anti-RNF213 antibodies were purchased from Sigma-Aldrich (Cat# HPA003347). Mouse monoclonal anti-SAG1 (clone DG52) antibodies were described in(51). Rabbit polyclonal anti-GRA7(52) and anti-SAG1 antibodies were kindly provided by Dr. John C. Boothroyd. HRP-conjugated Goat anti-Mouse/Rabbit/Rat IgG secondary antibodies were purchased from Jackson ImmunoResearch Laboratories Inc. (Cat# 111-035-003/112-035-003/115-035-003). HRP-conjugated Mouse anti-Goat IgG secondary antibodies were purchased from Santa Cruz Biotechnology (Cat# sc-2354). Goat anti-Mouse IgG (Alexa Fluor 448/594, Cat# A11029/A11032), Goat anti-Rat IgG (Alexa Fluor 594, Cat#A11007), and Goat anti-Rabbit (Alexa Fluor 488/594, Cat#A11008/A11037) secondary antibodies were purchased from Thermo Scientific.

### Animals

Six to eight-week-old female LEW rats (LEW/Crl; Strain Code: 004) and BN rats (BN/Crl; Strain Code: 091) were purchased from Charles River Laboratories. The rats were housed under pathogen-specific free conditions at the University of California, Davis animal facility and were allowed to acclimatize in the vivarium for at least a week undisturbed. In the facility, rats were housed in ventilated cages on corn bedding and provided with water and chow ad libitum. Cages were all on one rack at a housing density of three rats per cage. The rat housing room was on a 12-hours light/12-hours dark cycle with the temperature maintained at 22-25°C and the humidity range of 30-70%. The rats were monitored twice daily by veterinarians, and cage bedding was changed every two weeks. All animal experiments were performed in strict accordance with the recommendations in the Guide for the Care and Use of Laboratory Animals of the National Institutes of Health and the Animal Welfare Act, approved by the Institutional Animal Care and Use Committee at the University of California, Davis (Assurance Number: A-3433-01).

### Culture of cells and parasites

Human foreskin fibroblasts (HFFs, gift from Dr. John C. Boothroyd) were cultured in Dulbecco’s modified Eagle’s medium (DMEM) containing 10% fetal bovine serum (FBS), 2 mM L-glutamine, 100 U/mL penicillin/streptomycin, and 10 μg/mL gentamicin. Primary rat bone marrow-derived macrophages (BMDMs) were obtained by differentiating and cultivating bone marrow cells isolated from the tibia and femur of LEW rats or BN rats in Macrophage Differentiation Media (DMEM containing 20% FBS, 2 mM L-glutamine, 10 mM HEPES, 1 x non-essential amino acids, 1 mM sodium pyruvate, 100 U/mL penicillin/streptomycin, 10 μg/mL gentamicin, and 30% L929 conditioned medium) for 7 days. Fully differentiated rat BMDMs used for parasite infection or other experiments were cultured in Complete Macrophage Media (DMEM containing 10% FBS, 2 mM L-glutamine, 10 mM HEPES, 1 x non-essential amino acids, 1 mM sodium pyruvate, 100 U/mL penicillin/streptomycin, 10 μg/mL gentamicin, and 30% L929 conditioned medium). Lenti-X cells and HEK293T cells were DMEM containing 10% FBS, 2 mM L-glutamine, 10 mM HEPES, 1 x non-essential amino acids, 1 mM sodium pyruvate, 100 U/mL penicillin/streptomycin, and 10 μg/mL gentamicin.

SF9 insect cells were maintained in HyClone SFX-Insect Cell Media (Cytiva) supplemented with 1X antibiotic-antimycotic (Thermo Scientific, Cat# 15240096) at 27°C with constant shaking at 100 RPM. SF9 cells were recently purchased from the manufacturer and were not authenticated, and these cells were not regularly tested for mycoplasma contamination.

*Toxoplasma gondii* strains RH-Cas9(53), RHΔ*ku80*Δ*hxgprt*(54), RHΔ*hxgprt*(28), RHΔ*gra35*(28), RHΔ*gra35* + *GRA35HA*(28), RHΔ*ku80*Δ*hxgprt*-*GRA23-HA-FLAG::DHFR*(22), and RH-GRA15_II_-HA(25) were routinely passaged *in vitro* on monolayers of HFFs at 37 °C in 5% CO2. All cells and parasite strains were tested negative for mycoplasma contamination by PCR.

### Plasmid construction

All the plasmids and primers used in this study are listed in Table S2. The plasmid for making the GRA35 endogenously HA-tagged *Toxoplasma* strain was generated by amplifying and inserting ∼1.6 kb upstream of the stop codon of GRA35 into pLIC-3xHA::DHFR vector using ligation-independent cloning (54). To ectopically express C-terminal HA-tagged GRA6, GRA42, or GRA43 in mammalian cells, the coding sequence of mature GRA6, GRA42, or GRA43 (without signal peptide) was PCR amplified using primers listed in Table S2 and flanked with the HA epitope coding sequence before the stop codon followed by cloning into pcDNA3.1+ (Thermo Scientific, Cat# V79020) between *KpnI* and *EcoRI* sites. The mammalian expression vector containing C-terminal HA-tagged GRA35 without signal peptide named pcDNA3.1-GRA35-HA was constructed in our previous study(28). To construct the plasmid expressing the GRA35 C-terminus (after the transmembrane domain), the coding sequence of GRA35^142aa-378aa^ was PCR amplified using primers listed in Table S2 and flanked with the HA epitope coding sequence before the stop codon followed by cloning into pcDNA3.1+ between *KpnI* and *EcoRI* sites. To generate the mammalian expression construct containing full-length rat ITCH, the coding sequence of rat ITCH (Accession Number: XM_008762336) was amplified using primers listed in Table S2 and cloned into *SrfI* and *EcoRI*-linearized pCMVtag2B vector (N-terminal FLAG epitope-containing plasmid from Agilent Technologies, Cat# 211172) using Gibson Assembly (New England Biolabs, Cat# E5510S). The other pCMVtag2B vectors containing different truncated versions of rat ITCH (**Fig 2F**) were generated via Q5 Site-Directed Mutagenesis Kit (New England Biolabs, Cat#E0554S) by circularizing the PCR products amplified with the primers listed in Table S2. The DPP9 expression vector was generated by cloning the coding sequence of full-length rat DPP9 (Accession Number: NM_001305241) flanked with the N-terminal V5 epitope tag sequence into pcDNA3.1+ plasmid between *KpnI* and *EcoRI* sites. To generate SF9 protein expression vectors, the coding sequence of LEW rat NLRP1, BN rat NLRP1 (Accession Number: HM_060628), or rat DPP9 was subcloned into pFastBac HTB with a C-terminal FLAG epitope (named His-TEV-LEW-rNLRP1-FLAG, His-TEV-BN-rNLRP1-FLAG, or His-TEV-rDPP9-FLAG). To generate the Lentiviral expression construct containing LEW rat NLRP1, the coding sequence of LEW rat NLRP1 (Accession Number: HM_060633) was amplified and flanked with an EGFP tag at the N-terminus and an MYC epitope at the C-terminus before the stop codon followed by cloning into EcoRV-linearized pLenti-CMV-Puro-DEST plasmid (Addgene #17452) using Gibson Assembly.

### Genome-wide CRISPR/Cas9 screen in *Toxoplasma*

To generate a genome-wide knockout parasite population, 500 μg of AseI-linearized sgRNA library, which is a mixture of pU6-DHFR plasmids containing 10 different sgRNAs against each of the 8156 *Toxoplasma* genes(53), were transfected into 5 x 10^8^ of RH-Cas9 parasites (100 μg of library plasmid for each 1 x 10^8^ of parasites per transfection) followed by infection of HFFs at an MOI = 0.5. After cultivating in DMEM containing 1% FBS, 2 mM L-glutamine, 100 U/mL penicillin/streptomycin, 10 μg/mL gentamicin, and 40 μM chloramphenicol (CAT) (Sigma-Aldrich, Cat# C0378-5) for 24 h, the transfected parasites were grown in the DMEM containing 10% FBS, 2 mM L-glutamine, 10 mM HEPES, 1 x non-essential amino acids, 1 mM sodium pyruvate, 100U/mL penicillin/streptomycin, 10 μg/mL gentamicin, 40 μM CAT, 1 μM pyrimethamine, and 10 μg/mL DNase I (New England Biolabs, Cat#M0303S) for continuous passages in HFFs. To screen the parasite mutants that do not activate the NLRP1 inflammasome, LEW rat BMDMs were infected with the mutant pool consistent with at least 1×10^7^ of parasites harvested from the 4th lytic cycle (screen #1) or 1st lytic cycle (screen #2) in HFFs at an MOI = 0.2 for 2h. After washing out the extracellular parasites with PBS, the medium was replaced with a medium containing 30 mg/ml dextran sulfate to block the reinvasion of parasites released from pyroptotic macrophages. At 24 h post-infection, extracellular parasites lysed from pyroptotic cells were removed by washing with PBS for 3 times. The surviving cells containing parasite mutants unable to activate the NLRP1 inflammasome were collected and seeded onto a monolayer of HFFs. To maintain the parasite mutant diversity, 10% of the parasite population lysed from HFFs were passaged to the next round selection in LEW rat BMDMs. After 4 rounds of selection, we extracted genomic DNA from 1 x 10^7^ parasites and used it to amplify the sgRNAs with a barcoding primer via PCR. The resulting sample was then submitted for Illumina sequencing at the Genome Center of the University of California, Davis using a NextSeq (Illumina) with single-end reads using primers (P150 and P151) listed in Table S2.

The analysis of the library screen data was performed in R (www.R-project.org) version 4.2.0, Excel (Microsoft Office) version 16.72, and previously described custom software(53, 55). Briefly, the raw reads for each sgRNA were determined by aligning Illumina sequencing data to the sgRNA sequences presented in the library and counting the number of exact matches. To identify the genes that underwent positive selection, the raw reads of each sgRNA after 4th round selection in LEW rat BMDMs were compared to the library input, and the positive selection *p*-value of each gene was calculated using the MAGeCK algorithm(56). Genes were considered as high-confident candidates if they had at least 3 positively enriched sgRNAs in 4th round selection *vs.* library input and met a significance threshold of positive selection *p* < 0.05 in both screens.

### Generation of parasite strains

To generate a *Toxoplasma* strain expressing C-terminal HA-tagged GRA35, RHΔ*ku80*Δ*hxgprt* parasites were transfected with the plasmid pLIC-GRA35-3xHA-DHFR followed by selection with 3 μM pyrimethamine. After cloning by limiting dilution, the presence of *GRA35-3xHA* in the parasites was determined by immunofluorescence assays.

### Generation of NLRP1-expressing HEK293T cells

HEK293T cells stably expressing LEW rat NLRP1 were generated using the Lentiviral expression system. Lentiviral vector pLenti-CMV-Puro-DEST containing EGFP-rNLRP1-MYC was transfected into Lenti-X cells together with packaging plasmid psPAX2 (Addgene #12260) and Lentiviral envelope plasmid pMD2.G (Addgene #12259) using X-tremeGENE 9 DNA transfection reagent (Sigma-Aldrich, Cat# 6365787001) according to the manufacturer’s instructions. At 48 h post-transfection, virus-containing culture supernatant was collected, followed by mixing with polybrene (Sigma-Aldrich, Cat# TR-1003-G) at a final concentration of 8 μg/mL and adding into 6-well plates containing ∼50% confluent HEK293T cells. After 24 h, the cells were selected with puromycin (Sigma-Aldrich, Cat# 540411) at a concentration of 10 μg/mL for 72 h. After cloning by limiting dilution in 96-well plates (∼ one cell per well) with puromycin selection, the positive clones of NLRP1-expressing HEK293T cells were verified by the expression of GFP and MYC using immunoblotting.

### CRISPR/Cas9-mediated gene deletion in primary rat BMDMs and HFFs

CRISPR/Cas9 technology was used to generate an ITCH or NLRP1 deletion in LEW rat BMDMs. Three sgRNAs targeting the beginning of exon 2 of the rat *Itch* gene and two sgRNAs targeting the first exon of the rat *Nlrp1* gene were designed using Synthego’s online Knockout Guide Design tools (https://design.synthego.com/<u>#/</u>) and synthesized by Synthego (Redwood City, CA) with modification of 2’-O-methyl at the 3 first and last bases and 3’ phosphorothioate internucleotide linkages at the first three 5’ and 3’ terminal RNA residues with the purpose to improve the editing efficiency and minimize nucleotide acid-induced innate immune response in macrophages. As a scrambled control, sgRNAs targeting *E.coli* LacZ (Accession Number: NP_414878) were chosen and synthesized. All the sgRNA sequences are listed in Table S2. The *S. pyogenes* Cas9-NLS purified protein was obtained from QB3 Macrolab at the University of California, Berkeley. To make gene deletions, 4×10^6^ Lewis rat BMDMs were electroporated with *in vitro* assembled CRISPR/Cas9 ribonucleoprotein (400 pmol of Cas9-NLS protein combined with 600 pmol of sgRNAs) using the Neon transfection system (Thermo Scientific, Cat #5000S) with the following program: 1680 Volts/ 20 ms/1 pulse. Cells electroporated with only Cas9-NLS protein without sgRNAs were used as a negative control. After recovery in Complete Macrophage Media for 48 h, 1×10^5^ cells were used for genomic DNA isolation while other cells were seeded onto 96 well plates or coverslips (1×10^5^ cells per well or coverslip) for cell viability assay and parasite per vacuole counting. To check the editing efficiency, the CRISPR/Cas9 editing region was amplified from the *Itch* or *Nlrp1* genomic locus using primers listed in Table S2 followed by Sanger sequencing of PCR products and analysis with Inference of CRISPR Edits provided by Synthego (https://ice.synthego.com/#/).

To knockout *Itch* in HFFs, ready-to-use lentivirus particles containing three unique sgRNA targeting human *Itch* or viruses containing scrambled sgRNA were purchased from Applied Biological Materials Inc. (Richmond, BC, Canada). These Lentiviral particles were individually mixed with 8 μg/mL of polybrene and transduced into 6-well plates containing ∼50% confluent HFFs at the MOI of 5. After 24 h, the medium was replaced with fresh culture medium containing 1.5 μg/mL of puromycin and the HFFs were cultured for 5∼7 days (with a change of puromycin medium every other day) to select the stably transduced cells. Once the cells became ∼80% confluent, the HFF populations were expanded in T25 culture flasks, followed by checking for the knockout of ITCH using immunoblotting.

### Cell viability measurement using MTS and LDH assays

To determine the cell viability of rat BMDMs, a previously described MTS assay was performed(28). Briefly, 1 x 10^5^ of BMDMs isolated from LEW rats or BN rats were seeded into one well of 96-well plates followed by *Toxoplasma* infection at an MOI of 1 or VbP treatment at a concentration of 2 µM. After 24 h infection/treatment, cell viability was measured by adding 3 - (4,5 - dimethylthiazol - 2 - yl) - 5 - (3 - carboxymethoxyphenyl) - 2 - (4 - sulfophenyl) - 2H - tetrazolium (MTS) into culture media followed by reading the OD_490_ value after 1.5 h. Raw absorbance of cells without infection/treatment was considered as 100 percent, whereas 0 was used to stand for cells treated with lysis buffer before adding MTS. The percentage of viable cells was calculated by expression of relative absorbance of *Toxoplasma*-infected or VbP-treated cells vs. cells without infection/treatment.

To measure the cell viability of HFFs, a previously described LDH assay was performed(57). Briefly, 2×10^4^ of HFFs seeded into one well of 96-well plates were stimulated with 10U/mL human IFNγ or left unstimulated for 24 h followed by the infection with different *Toxoplasma* strains at MOI = 1. At 24 h post-infection, 100 µL of culture supernatant was mixed with LDH reagent (Sigma-Aldrich, Cat# 11644793001) followed by reading the OD490 value after 20 min incubation. Raw absorbance of cells treated with 2% Triton X-100 (lysis control) was considered as 100 percent of LDH release, whereas 0 was used for the uninfected cells treated with lysis buffer before measuring LDH release. The percentage of LDH release in parasite-infected cells was calculated using the formula: %LDH release = (OD_490_ value of infected cells – OD_490_ value of uninfected cells)/(OD_490_ value of lysis control – OD_490_ value of uninfected cells).

### Parasite per vacuole counting

To count parasites per vacuole in LEW rat BMDMs, coverslips containing 2 x 10^5^ LEW rat BMDMs were infected with 1 x 10^5^ parasites for 30 min followed by removing the uninvaded parasites by washing 3 times with PBS. After 24 h infection, the cells were fixed with 4% Paraformaldehyde (PFA) for 20 min followed by permeabilization/blocking with PBS containing 3% (w/v) BSA, 5% (v/v) goat serum (Thermo Scientific, Cat# 16210072), and 0.1% Triton X-100 for 30 min. The parasites were detected by incubating the coverslips with mouse anti-SAG1 DG52 (1:100 dilution) and rabbit anti-GRA7 (1:3000 dilution) antibodies for 1 h at room temperature. After incubating with secondary antibodies Alexa Fluor 488-conjugated goat anti-mouse IgG (1:3000 dilution) and Alexa Fluor 594-conjugated goat anti-rabbit IgG (1:3000 dilution) together with DAPI (Sigma-Aldrich, Cat# D9542) at the final concentration of 1 μg/mL, the coverslips were mounted with Vecta-Shield mounting oil and the microscopy was performed with NIS-Elements software (Nikon) and a digital camera (CoolSNAP EZ; Roper Scientific) connected to an inverted fluorescence microscope (Eclipse Ti-S; Nikon). The number of parasites in at least 100 vacuoles was observed, counted, and quantified.

### Invasion assay

The invasion assay was performed as previously described(58) with minor modification. Briefly, 1×10^5^ LEW rat BMDMs seeded onto coverslips were treated with 0.5 μM MG132 or 30 mg/mL dextran sulfate (as a non-invasion control) for 2 h followed by infection with 2×10^5^ RHΔ*hxgprt* parasites for another 30 min. After washing with PBS three times, the cells were fixed with 4% PFA for 20 min at room temperature and blocked with PBS containing 3% (w/v) BSA for 30 min at room temperature, and extracellular parasites were stained with rabbit anti-SAG1 (1:5000 dilution in PBS containing 3% BSA) for 1 h at room temperature. The cells were then permeabilized in PBS containing 3% (w/v) BSA, 5% (v/v) goat serum, and 0.1% Triton X-100 for 30 min at room temperature followed by incubation with mouse monoclonal anti-SAG1 DG52 (1:100 dilution) for 1 h at room temperature. Alexa Fluor 594-conjugated goat anti-rabbit IgG (1:3000 dilution) and Alexa Fluor 488-conjugated goat anti-mouse IgG (1:3000 dilution) were used as secondary antibodies, while 1 μg/mL of DAPI was added to the secondary antibody solution to stain host nuclei. The coverslips were mounted with Vecta-Shield mounting oil, and the microscopy was performed with NIS-Elements software (Nikon) and a digital camera (CoolSNAP EZ; Roper Scientific) connected to an inverted fluorescence microscope (Eclipse Ti-S; Nikon). To determine the parasite invasion efficiency, at least 10 random fields were observed for all samples, and the total number of green/yellow parasites (intracellular + extracellular) and red parasites (extracellular) were counted and used to calculate the ratio of intracellular parasites (number of green/yellow parasites subtract the number of red parasites) *vs.* host nucleus.

### Selective permeabilization

The selective permeabilization was performed as previously described(59). Briefly, HFFs grown on coverslips were infected with RHΔ*ku80*Δ*hxgprt*-*GRA35-3xHA::DHFR* (endogenously HA-tagged GRA35-expressing strain) for 20 h. After fixation in 4% PFA for 10 min at room temperature, the cells were quenched with PBS containing 100 mM glycine for 5 min at room temperature followed by semi-permeablization with PBS containing 0.001% digitonin for 5 min at 4°C. The samples were then incubated with blocking buffer (PBS containing 10% FBS) for 30 min at room temperature, probed with antibodies against the HA epitope (1:500 dilution in blocking buffer) together with antibodies against SAG1 (1:5000 dilution in blocking buffer) for 1h at room temperature. After incubating with secondary antibodies Alexa Fluor 594-conjugated goat anti-rat IgG (1:3000 dilution in blocking buffer) and Alexa Fluor 488-conjugated goat anti-rabbit IgG (1:3000 dilution in blocking buffer) together with 1 μg/mL of DAPI, the coverslips were mounted with Vecta-Shield mounting oil and the microscopy was performed with NIS-Elements software (Nikon) and a digital camera (CoolSNAP EZ; Roper Scientific) connected to an inverted fluorescence microscope (Eclipse Ti-S; Nikon).

### Immunofluorescence assay for detecting the recruitment of ITCH, ubiquitin, and RNF213 to the PVM

Scrambled control or *Itch*-knockout HFFs grown on coverslips were stimulated with 10U/mL human IFNγ or left unstimulated for 24 h, and subsequently infected with GFP-expressing *Toxoplasma* strains (wild-type or Δ*gra35*) at an MOI = 1 for another 4 h. After fixation in 4% PFA for 10 min at room temperature, the cells were permeabilized/blocked with PBS containing 3% (w/v) BSA, 5% (v/v) goat serum, and 0.1% Triton X-100 for 30 min followed by incubating with mouse anti-ITCH (1:100 dilution), mouse anti-Ubiquitin (1:250 dilution), or rabbit anti-RNF213 (1:400 dilution) for 1 h at room temperature or overnight at 4°C. After incubating with secondary antibodies Alexa Fluor 594-conjugated goat anti-mouse IgG or goat anti-rabbit IgG (1:3000 dilution) together with 1 μg/mL of DAPI, the coverslips were mounted with Vecta-Shield mounting oil and the microscopy was performed with NIS-Elements software (Nikon) and a digital camera (CoolSNAP EZ; Roper Scientific) connected to an inverted fluorescence microscope (Eclipse Ti-S; Nikon). To quantify the percentage of recruitment, at least 150 vacuoles were observed and counted.

### Immunoprecipitation

To identify the host interaction partner of GRA35, two independent immunoprecipitations were performed in *Toxoplasma*-infected rat BMDMs. In the first experiment, 4×10^7^ of LEW rat BMDMs were treated with 50 mM VX-765 (Caspase-1/11 inhibitor) for 2 h followed by infection with RHΔ*ku80*Δ*hxgprt*-*GRA35-3xHA::DHFR* or RHΔ*ku80*Δ*hxgprt*-*GRA23-HA-FLAG::DHFR* (as a control) at an MOI of 3. After 6 h infection, the cells were harvested and lysed in 1 mL of IP-lysis buffer A (125 mM Tris-Cl pH7.5, 150 mM NaCl, 1% NP40) containing 1x Halt^TM^ protease and phosphatase inhibitor and 1 mM phenylmethylsulfonyl fluoride (PMSF) for 30 min on ice. The lysate was centrifuged for 30 min at 18,000 x g, 4°C, and the supernatant (soluble fraction) was incubated with 20 μL of anti-HA magnetic beads for 3 h at 4°C with rotation. The beads were washed and resuspended in IP-lysis buffer A followed by Mass Spectrometry analysis. For the 2nd independent experiment, 2×10^8^ BMDMs isolated from LEW rats or BN rats were pre-treated with 50 mM VX-765 for 2h followed by infection with RHΔ*ku80*Δ*hxgprt*-*GRA35-3xHA::DHFR* or RH-GRA15_II_-HA (as a control) at an MOI of 3 for another 8 h. The cells were lysed in 4 mL of IP-lysis buffer B (25 mM HEPES, 150 mM NaCl, 1 mM EDTA, 1 mM EGTA, 0.65% NP40) containing 1x Halt^TM^ protease and phosphatase inhibitor cocktail and 1 mM PMSF for 30 min on ice. The lysate was centrifuged for 30 min at 18,000 x g, 4°C, and the supernatant was incubated with 100 μL anti-HA magnetic beads at 4°C overnight with rotation. After washing, the beads resuspended in IP-lysis buffer B were subjected to Mass Spectrometry analysis.

To confirm the interaction between GRA35 and ITCH, HEK293T cells were transfected with FLAG-ITCH expressing plasmid pCMVtag2B-ITCH together with GRA35-HA expressing plasmid pcDNA3.1-GRA35HA at a 1:1 ration using X-tremeGENE 9 DNA transfection reagent according to the manufacturer’s instructions. As controls, cells were also co-transfected with pCMVtag2B-ITCH and pcDNA3.1 vector containing the coding sequence of other dense granule proteins (pcDNA3.1-GRA42HA, pcDNA3.1-GRA43HA, or pcDNA3.1-GRA6HA). To check the interaction between the GRA35 C-terminus and ITCH, HEK293T cells were transfected with pCMVtag2B-ITCH together with pcDNA3.1 vector expressing HA-tagged GRA35 C-terminus or full-length GRA35 (as a control). To determine the ITCH domain interacting with *Toxoplasma* GRA35, the pcDNA3.1 vector expressing the HA-tagged GRA35 C-terminus was co-transfected with pCMVtag2B containing full-length or truncated versions of ITCH into HEK293T cells. After 30 h, transfected cells were scraped in ice-cold PBS and lysed in IP-lysis buffer B containing 1x Halt^TM^ protease and phosphatase inhibitor cocktail and 1 mM PMSF for 30 min on ice. The lysate was centrifuged for 30 mins at 18,000 x g, 4°C, and the supernatant was incubated with anti-HA magnetic beads at 4°C overnight with rotation. After washing with IP-lysis buffer B for three times, proteins bound to the beads were solubilized in SDS loading buffer by boiling for 5 min and examined by immunoblotting analysis. To perform reciprocal IP, HEK293T cell transfected with HA-tagged GRA35 C-terminus together with pCMVtag2B-ITCH or pCMVtag2B empty vector (as a control) were lysed in IP-lysis buffer B containing 1x Halt^TM^ protease and phosphatase inhibitor cocktail and 1 mM PMSF after 30 h of transfection followed by immunoprecipitation with anti-FLAG Magnetic Agarose (Thermo Scientific, Cat # A36797) at 4°C overnight with rotation. After washing with IP-lysis buffer B for three times, proteins bound to the agarose were solubilized in SDS loading buffer by boiling for 5 min and examined by immunoblotting analysis.

### Mass spectrum analysis

To identify proteins in GRA35-immunoprecipitated samples, the magnetic beads after immunoprecipitation were sent to the Proteomics Core Facility of the University of California, Davis, for mass spectrometry analysis. After overnight on-bead digestion with trypsin, the peptide extracts were analyzed by LC-MS/MS using a Thermo Scientific Q Exactive Plus Orbitrap Mass Spectrometer in conjunction with Thermo Scientific Proxeon Easy-nLC II HPLC and Proxeon nanospray source (for 1st independent experiment) or using a Thermo Scientific Dionex UltiMate 3000 RSLC system in conjunction with Thermo Scientific Orbitrap Exploris 480 instrument (for 2nd independent experiment). Mass spectrometry raw files were searched using Fragpipe 16.0(60) against the UniProt *Toxoplasma gondii* and rat database with default search settings. Decoy sequences were generated, appended and laboratory contaminates added within Fragpipe. Decoy False Discovery Rates were controlled at 1% maximum using both the Peptide and Protein prophet algorithms(61). Search results were loaded into Scaffold (version Scaffold_5.0.1, Proteome Software Inc., Portland, OR) for visualization purposes. Proteins that contained similar peptides and could not be differentiated based on MS/MS analysis alone were grouped to satisfy the principles of parsimony. Proteins sharing significant peptide evidence were grouped into clusters. The total unique spectrum count for all samples is available in Table S3.

### Protein expression

Recombinant His-TEV-rDPP9-FLAG, His-TEV-Lew-rNLRP1-FLAG, and His-TEV-BN-rNLRP1-FLAG were purified similarly to human His-DPP9 (https://www.protocols.io/groups/hao-wu-lab). Baculoviruses containing each of these proteins were prepared using the Bac-to-Bac system (Invitrogen) and used to generate baculovirus-infected SF9 insect cells. To express His-TEV-rDPP9-FLAG, His-TEV-Lew-rNLRP1-FLAG, or His-TEV-BN-rNLRP1-FLAG, 1 mL of the corresponding baculovirus-containing cells was used to infect each L of SF9 cells. Cells were harvested 48 h after infection by centrifugation (1,682 x g, 20 min), washed once with phosphate-buffered saline (PBS), flash-frozen in liquid nitrogen, and stored at -80 °C. The thawed pellet from 2 L of cells was resuspended in lysis buffer (80 mL, 25 mM Tris-HCl pH 8.0, 150 mM NaCl, 1 mM tris(2-carboxyethyl)phosphine abbreviated as TCEP, 5 mM imidazole), sonicated (3 s on 7 s off, 3.5 min total on, 50% power, Branson), and ultracentrifuged (186,000 x g, 1.5 h, 45 Ti fixed-angle rotor, Beckman). After centrifugation, the supernatant was incubated with 1 mL Ni-NTA resin at 4 °C for 1 h. The bound Ni-NTA beads were washed once in batch and subsequently by gravity flow using 50-100 CV wash buffer (25 mM Tris-HCl pH 8.0, 150 mM NaCl, 1 mM TCEP, 25 mM imidazole). The protein was eluted with buffer containing 500 mM imidazole (5 mL), spin concentrated to 0.5 mL (Amicon Ultra, 50 kDa MW cutoff), and further purified by size exclusion chromatography (25 mM Tris-HCl pH 7.5, 150 mM NaCl, 1 mM TCEP) on a Superdex 200 10/300 GL column (Cytiva). Peak fractions were pooled, aliquoted, and flash-frozen in liquid nitrogen for use in ubiquitination assays (below).

### *In vitro* ubiquitination assays

ITCH ubiquitination reactions were carried out using the manufacturer’s protocol for the Human ITCH/AIP4 Ubiquitin Ligase Kit (R&D Systems, K-270). 10X reaction buffer, 10X E1 enzyme, 10X E2 enzyme (UBE2L3), 10X ITCH E3 ligase, and 10X ubiquitin solution were combined on ice and supplemented with 1 μM rDPP9-FLAG, 1 μM Lew-rNLRP1-FLAG, 1 μM BN-rNLRP1-FLAG, or no substrate control (15 μL final volume per reaction). Ubiquitination was initiated with the addition of 10X Mg2+-ATP or water (negative control) to the reaction solution. Ubiquitination reactions were incubated at 37 °C for 2 h and then quenched with the addition of 4X SDS sample buffer.

### Immunoblotting

To detect protein interaction, total proteins from the immunoprecipitated samples were loaded and run onto 12% SDS-PAGE gels followed by transferring to a polyvinylidene difluoride (PVDF) membrane. To check the ITCH knockout efficiency, the lysis from at least 1×10^6^ of scrambled control HFFs or *Itch*-knockout HFFs (three independent clones) were loaded and run onto 12% SDS-PAGE gels followed by transferring to a PVDF membrane. Membranes were blocked in 5% milk in TBS supplemented with 1% Tween-20 (TBS-T) for 1 h at room temperature followed by incubation with primary antibodies diluted in the blocking buffer at 4°C overnight. After incubation with HRP-conjugated secondary antibodies, the protein of interest on the membranes was visualized using ProSignal® Femto ECL Reagent (Genesee Scientific, Cat# 20-302), and the images were acquired using KwikQuant Imager (Kindle Biosciences, LLC).

For *in vitro* ubiquitination, samples were run on either 4 to 15% or 4 to 20% Mini-PROTEAN TGX™ Tris-Glycine gels (BioRad) for 40-60 min at 160 V. Gels were transferred to nitrocellulose with the iBlot 2 Transfer System (Thermo Scientific). Membranes were blocked with phosphate buffered saline with 0.05% tween-20 (PBST) supplemented with 5% milk (PBST-M) for 60 min at ambient temperature, prior to incubating with primary antibody (in PBST-M) overnight at 4 °C. Blots were washed 3 times with PBST prior to incubating with secondary antibody (in PBST-M) for 60 min at ambient temperature. Blots were again washed 3 times and subsequently imaged with a BioRad ChemiDoc using ECL Western Blotting Substrate (Thermo Scientific 32106). Antibodies used include: Ubiquitin mouse monoclonal Ab (1:2000, Santa Cruz Biotechnology, sc-8017), ITCH goat Ab (1:2000, R&D Systems, K-270 proprietary), anti-FLAG-HRP mouse monoclonal Ab (1:5000, Sigma, A8592), anti-mouse IgG-HRP (1:10000, Cell Signalling Technology, 7076P2), and anti-goat IgG-HRP (1:10000, Santa Cruz Biotechnology, sc-2354).

### Plaque assay

Scrambled control or *Itch*-knockout HFFs seeded into 24-well plates were stimulated with human IFNγ (10 or 20 U/mL) or left unstimulated for 24 h followed by infection of 100 parasites into each well. Infected plates were incubated for 5 days at 37°C, and the number of plaques formed by parasite infection was observed and counted under the microscope using 4 x objective. To calculate the percentage of plaque loss, the following formula was used: [(Number of plaques in unstimulated HFFs - Number of plaques in IFNγ-stimulated HFFs)/Number of plaques in unstimulated HFFs] × 100. All experiments were performed at least 3 times with duplicate wells for each condition.

### Statistical analysis

All statistical analyses were performed using Prism (GraphPad) version 9.5. All the data are presented as mean ± standard deviation (SD), and the exact n values are mentioned in the figure legends. For all the calculations, *p* < 0.05 are considered a significant difference. To compare parasite growth of Δ*gra35* vs. wild-type parasites in IFNγ-activated HFFs, paired t-test was used. For the data with more than two groups with one variable, One-way ANOVA with Tukey’s multiple comparisons test was used. For one variable test with two groups, the two-way ANOVA with Tukey’s multiple comparisons test was used.

### Data Availability

The authors declare that all data supporting the findings of this study are available within the article and Expanded View Information files, including Source Data and uncropped immunoblotting images. CRISPR screen data including raw sequencing read counts are available in Table S1. Mass spectrometry data are available in Table S3. All unique materials (e.g., the variety of parasite and host cell lines described in this study) are available from the corresponding author (contact: Yifan Wang; Jeroen P.J. Saeij) upon reasonable request.

## ACKNOWLEDGMENTS

This study was supported by the National Institutes of Health (R01-AI080621 and R21-A151081 awarded to J.P.J.S., R01-AI144149 awarded to B.H.P., and R01-AI124491 awarded to H.W.) and NIH Pathway to Independence Award (R00-AI163285) to Y.W. We thank R.E. Vance for sharing the C-terminal NLRP1 (2A12) antibodies. We thank S. Lourido for sharing the sgRNA library and providing technical support for data analysis.

## CONFLICT OF INTERESTS

Dr. Hao Wu is a co-founder of Ventus Therapeutics. The other authors declare that they have no conflict of interest.

